# Generative AI-assisted Virtual Screening Pipeline for Generalizable and Efficient Drug Repurposing

**DOI:** 10.1101/2024.12.07.627340

**Authors:** Phuc Pham, Viet Thanh Duy Nguyen, Kyu Hong Cho, Truong Son Hy

## Abstract

Drug repurposing presents a valuable strategy to expedite drug discovery by identifying new therapeutic uses for existing compounds, especially for diseases with limited treatment options. We propose a Generative AI-assisted Virtual Screening Pipeline that combines generative modeling, binding pocket prediction, and similarity-based searches within drug databases to achieve a generalizable and efficient approach to drug repurposing. Our pipeline enables blind screening of any protein target without requiring prior structural or functional knowledge, allowing it to adapt to a wide range of diseases, including emerging health threats and novel targets where information is scarce. By rapidly generating potential ligands and efficiently identifying and ranking drug candidates, our approach accelerates the drug discovery process, broadening the scope and impact of repurposing efforts and offering new possibilities for therapeutic development. Detailed results and implementation can be accessed at https://github.com/HySonLab/DrugPipe.

## 1 Introduction

The discovery of new drugs is a costly and time-consuming process, often requiring substantial financial investment and many years of development. Traditional *de novo* drug discovery typically involves identifying a lead compound, followed by extensive preclinical and clinical testing, contributing to high costs and lengthy timelines. These challenges underscore the need for more efficient methods. Recent advancements in drug discovery have been driven by the integration of computational methods and artificial intelligence (AI), which have significantly accelerated the identification and optimization of potential therapeutic compounds, offering new avenues for treating various diseases. AI and computational tools have enhanced the speed and efficiency of drug discovery, reduced costs, improved target specificity, and provided valuable data-driven insights [1, 2]. Additionally, these advancements have supported the development of personalized medicine, enabling therapies tailored to individual genetic and molecular profiles [3]. However, challenges remain. One major issue is that while generative AI models can create ligands with high theoretical binding affinities and desirable pharmacological properties, these ligands may be difficult or even impossible to synthesize in practice. This challenge arises because generative AI models often prioritize certain chemical and biological criteria that do not always align with practical constraints in chemical synthesis and manufacturing, leading many promising AI-generated ligands to fail when transitioning from computational predictions to tangible, testable compounds.

Conversely, drug repurposing offers a promising alternative by discovering new therapeutic applications for existing drugs [4]. A key advantage of repurposing is the potential for substantial time and cost savings, as it involves compounds that have already been tested extensively for safety and efficacy, allowing for quicker transitions from research to clinical applications. This is especially crucial in urgent scenarios, such as pandemics, where the lengthy process of developing new drugs from scratch is impractical. Repurposing offers a faster response by leveraging the known safety and efficacy profiles of existing drugs to address emerging health threats. Furthermore, drug repurposing carries lower risk because it relies on drugs with well-documented pharmacokinetic and pharmacodynamic profiles, reducing the likelihood of unexpected adverse effects and increasing the chances of regulatory approval. However, drug repurposing faces significant challenges, such as the extensive time and resources needed to screen a vast number of existing drugs against new targets, making the process resource-intensive and time-consuming [5]. Additionally, many repurposing efforts have been limited to single diseases, restricting the broader applicability of findings and potentially missing opportunities to repurpose drugs for other conditions [6–9].

To address these challenges, we propose a Generative AI-assisted Virtual Screening Pipeline that enhances both efficiency and generalizability in identifying new therapeutic applications for existing drugs. Our approach integrates generative AI models for ligand generation, binding pocket prediction, and similarity-based searches within approved drug databases. By utilizing the speed of similarity-based searches, which are significantly faster than traditional docking simulations [10], our methodology rapidly generates and ranks potential drug candidates. Although docking simulations generally provide more precise insights into binding interactions, they are also computationally intensive and time-consuming. In contrast, the efficiency of our approach makes it well-suited for large-scale drug repurposing, where time and resource optimization are essential. Furthermore, the generalizability of our pipeline enables its application across a wide range of diseases, including scenarios with limited data, such as emerging health threats and novel targets. By prioritizing efficiency and adaptability, our framework offers a robust and versatile solution for accelerating the drug repurposing process.

To evaluate the generalizability and practical application of our pipeline, we conducted a comprehensive study using the DrugBank database, which includes protein targets and their corresponding approved drugs. This evaluation aimed to determine whether our pipeline could effectively identify real drugs in repurposing scenarios, such as COVID-19 and HIV. Given that our model relies on binding affinity, we hypothesize that the active compounds identified by the pipeline will likely represent effective drugs in real-world applications. However, recognizing that not all approved drugs exhibit the highest binding affinities due to factors like pharmacokinetics and safety profiles, we report the distribution of the position where the first approved drug for each target appears within our ranked list of predicted compounds. This analysis helps assess the pipeline’s ability to narrow down the search space for wet lab experiments to include real, approved drugs, thereby demonstrating the practical utility and broad applicability of our framework.

The contributions of this work are summarized as follows:

- We propose a novel Generative AI-assisted Virtual Screening Pipeline that leverages the strengths of generative AI models for efficient ligand generation and virtual screening, enhancing both the speed and accuracy of drug repurposing efforts.
- Our pipeline demonstrates high generalizability, as it can be applied across various diseases and targets without requiring prior knowledge of protein structures or known active compounds, making it especially valuable for emerging health threats and novel targets.
- We conduct a comprehensive evaluation using the DrugBank database, illustrating the pipeline’s effectiveness in identifying known drugs for critical diseases like COVID-19 and HIV. This includes an analysis of the pipeline’s performance in narrowing down the search space to prioritize real, approved drugs.

## 2 Related Work

### 2.1 Generative AI Models in Drug Discovery

Generative AI models have become powerful tools in drug discovery, enabling the design and optimization of novel molecular structures with specific desired properties [11–13]. Numerous studies have demonstrated the effectiveness of these models in generating promising drug candidates. For instance, many generative approaches have produced molecules by conditioning on specific protein binding pockets, resulting in high theoretical binding affinities and pharmacological properties [13–15]. These studies have shown substantial success in the design of ligands tailored to known protein targets. However, a significant limitation of these methods is their reliance on detailed structural information about the protein binding pocket. This dependency becomes a critical drawback in situations where such information is unavailable, as in the case of emerging pathogens like new viruses.

To address this limitation, in this work, we introduce an approach that integrates binding pocket prediction with generative AI models to enhance drug discovery in contexts where binding pocket details are not available. Our method first employs advanced binding pocket prediction techniques to identify multiple potential binding sites on target proteins. Subsequently, a generative AI model is used to design novel compounds optimized for each of these predicted pockets. This strategy not only enables the discovery of new drug candidates when binding pocket information is limited but also enhances the diversity of generated compounds by targeting a variety of potential binding sites, thereby increasing the chances of identifying effective therapeutic agents.

### 2.2 Current Approaches in Virtual Screening for Drug Repurposing

Virtual screening has become an essential tool in the drug repurposing process, particularly for the rapid identification of therapeutic candidates for emerging diseases. These approaches utilize computational methods to efficiently screen extensive compound libraries, providing significant advantages in terms of speed and cost-effectiveness compared to traditional drug discovery processes [16].

Despite promising outcomes, several limitations persist across existing research efforts. As illustrated in Table 1, many virtual screening and drug repurposing strategies depend on extensive prior knowledge about the protein target, such as detailed information on binding pockets, approved drugs, and known active compounds. This reliance can be a significant drawback, especially when dealing with novel or poorly characterized targets, where such data may be incomplete or unavailable. Additionally, a substantial portion of existing studies relies heavily on docking simulations using tools like AutoDock Vina [6, 17–19]. Although Vina is well-regarded for its accuracy, the process of docking compounds can vary significantly in duration. For simpler receptors and suitable ligands, docking might be completed in minutes. However, with complex protein structures or incompatible ligands, simulations can take several hours per compound. This extended time is due to the need for the algorithm to explore numerous potential binding conformations and interactions, especially when the ligand does not easily fit the binding site. Such variability makes large-scale docking simulations resource-intensive and can significantly slow down the drug discovery process when screening extensive compound libraries [20].

**Table 1:** Comparison of Virtual Screening for Drug Repurposing Approaches Based on Input and Output Characteristics.

| Approach | Input |  |  | Output |  |
| --- | --- | --- | --- | --- | --- |
|  | Protein Structure | Known Active Compounds | Binding Pocket | Drug Candidates | Binding Poses |
| Jang et al. [6] | ✓ | ✓ | ✓ | ✓ | ✓ |
| Morselli et al. [21] | ✓ | ✓ | × | ✓ | × |
| Chen et al. [18] | ✓ | × | ✓ | ✓ | ✓ |
| Gan et al. [17] | ✓ | × | × | ✓ | ✓ |
| Pirolli et al. [19] | ✓ | × | × | ✓ | ✓ |
| Ours | ✓ | × | × | ✓ | ✓ |

Many existing approaches are tailored to specific diseases [6, 19, 21, 22], limiting their generalizability and broader applicability. This narrow focus can hinder the discovery of repurposing opportunities across different therapeutic areas, reducing their overall impact. Moreover, such specificity poses significant challenges when addressing emerging health threats, as these approaches may struggle to adapt to novel pathogens. Rapid and flexible response capabilities are essential for addressing new diseases, where prior detailed knowledge of targets is often unavailable.

Given these limitations, there is a clear need for more versatile and efficient approaches in virtual screening and drug repurposing. Our work addresses these challenges by developing a generalizable framework that minimizes reliance on detailed prior knowledge of protein targets. By incorporating generative AI models for ligand design, the framework eliminates the need for known active compounds and reduces dependency on resource-intensive docking simulations, accelerating screening for large compound libraries. This flexibility, powered by generative AI, enables the framework to adapt to a wide range of diseases, including novel pathogens, ensuring responsiveness to diverse health scenarios.

## 3 Background

### 3.1 Graph neural networks

Graphs are versatile data structures capable of representing intricate relational data, consisting of nodes and edges. They are prevalent across various fields, including social networks [23], computational chemistry [24–26], biology [27], recommendation systems [28], chip design [29], code understanding [30], semi-supervised learning [31], and more.

#### Graph Representation

A graph *G* = (*V, E*) consists of a set of nodes *V* and edges *E*, where each node *v* ∈ *V* is associated with a feature vector *x*_*v*_, and each edge (*u, v*) ∈ *E* represents a relationship between two nodes *u* and *v*. The graph structure is represented by an adjacency matrix *A*, where *A*_*uv*_ = 1 if there is an edge between nodes *u* and *v*, and *A*_*uv*_ = 0 otherwise.

#### Message passing GNNs

The initial graph neural network (GNN) model introduced by [32, 33] was the pioneering deep learning approach for graph-structured data. It conceptualizes the task of graph embedding as a process of information diffusion, where nodes transmit information to their neighboring nodes, continuing until a stable equilibrium state is achieved. In each layer of a GNN, every node updates its feature vector by aggregating the information from its neighboring nodes. This is done through a message-passing mechanism, where messages are passed along the edges of the graph. The message-passing process typically involves two steps:

- **Aggregation**: For each node *v*, the feature vectors of its neighboring nodes 𝒩(*v*) are aggregated using a permutation-invariant function such as sum, mean, or max.
- **Update**: The aggregated message is then combined with the node’s current feature vector *h*_*v*_ to produce an updated representation. This can be done through a learnable transformation, such as a neural network, often in the form of a linear transformation followed by a non-linear activation function:

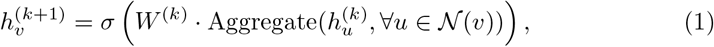

where 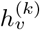 is the feature vector of node *v* at layer *k, W* ^(*k*)^ is a learnable weight matrix, *σ* is a non-linear activation function (e.g., ReLU [34]), and the aggregation function computes a summary of the neighboring nodes’ features.

### 3.2 Equivariance neural networks

Equivariant Neural Networks (ENN) are models designed to respect the symmetries present in the data, ensuring that certain transformations of the input result in predictable transformations of the output. This is particularly important in domains such as physics, chemistry, and computer vision, where data often exhibits intrinsic symmetries. In ENNs, these symmetries are encoded into the architecture, leading to models that generalize better, are more data-efficient, and respect physical laws. The two key types of equivariance discussed here are **permutation equivariance** [35] and **rotation equivariance** [36]. These correspond to different symmetries depending on the structure of the data.

#### Permutation Equivariance

Permutation equivariance arises when the data is unordered, such as in graphs or sets. For example, molecular structures can be viewed as graphs, where nodes represent atoms and edges represent bonds, and permuting the atoms should not change the physical properties of the molecule. Formally, a function *f* : *X* → *Y* is permutation-equivariant with respect to the symmetric group *S*_*n*_, which permutes the nodes of a graph. This means that for any permutation Π ∈ *S*_*n*_, the function satisfies the property:

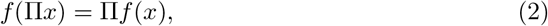

where Π is an *n* × *n* permutation matrix, and *x* ∈ ℝ^*n×d*^ is the input node feature matrix with *n* nodes and *d* features per node. This property ensures that if the input nodes are permuted, the output is permuted in the same way, making models like Graph Neural Networks (GNNs) suitable for problems with unordered data. In a GNN, node features *h*_*i*_ are updated by aggregating information from neighboring nodes:

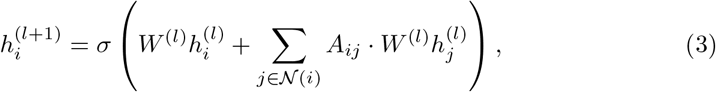

where 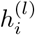 is the feature vector of node *i* at layer *l, A*_*ij*_ is the adjacency matrix of the graph, *W* ^(*l*)^ are the learnable weights, and *σ* is a non-linear activation function. This architecture respects permutation equivariance because permuting the nodes permutes both the features and the adjacency matrix, leaving the update rule unchanged.

#### Rotation Equivariance

Rotation equivariance is relevant when the data resides in physical space and is subject to transformations such as rotations and reflections. For instance, in molecular modeling, atomic coordinates can be rotated or reflected, but the physical properties of the molecule should remain consistent under these transformations. A function *f* is said to be rotation-equivariant if applying a rotation or reflection to the input *x* results in an equivalent transformation of the output:

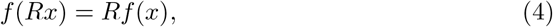

where *R* ∈ O(3) is an orthogonal matrix representing a rotation or reflection in three-dimensional space. Here, *x* ∈ ℝ^*n×*3^ represents the 3D coordinates of *n* points (e.g., atoms), and *f* (*x*) might represent quantities like forces or velocities that should transform in the same way under rotations.

A common example of a rotation-equivariant network is an E(3)-equivariant network, where the goal is to respect transformations from the Euclidean group *E*(3), which includes rotations, translations, and reflections. In these networks, both node features (such as atomic types) and geometric coordinates (3D positions) are updated while preserving equivariance.

For instance, consider a feature update rule in an E(3)-equivariant network:

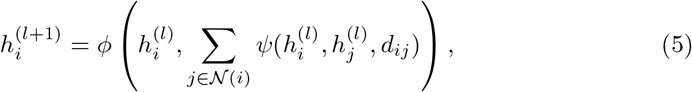

where *d*_*ij*_ = || *x*_*i*_ − *x*_*j*_ ||^2^ is the squared Euclidean distance between nodes *i* and *j*, and *ϕ, ψ* are learnable functions. For the coordinate updates, the equivariant transformation rule is applied:

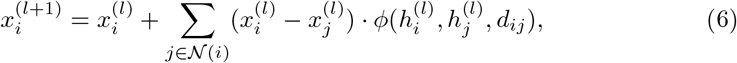

where the term 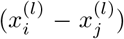 ensures that the update respects the relative distances between the points, making it equivariant under rotations and translations.

In diffusion models, equivariance can also be applied to the distribution of data over time. A conditional distribution *p*(*y*|*x*) is said to be equivariant to a group *G* (e.g., the Euclidean group *E*(3)) if it satisfies:

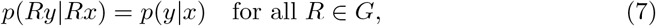

where *x* and *y* could represent the input and output distributions over time, and *R* is a transformation such as rotation. This ensures that the diffusion process respects the symmetries of the data at every time step. In practice, this is useful in denoising or generative models where the goal is to generate physically realistic 3D structures that respect rotational or reflection symmetries.

### 3.3 Protein representation learning

In recent years, protein representation learning has emerged as a key focus in structural biology such as property prediction [37, 38], affinity prediction [13, 39], protein design [40–43], and rigid docking [44]; and drug design [13, 45–47]. Traditional methods such as one-hot encoding or sequence-based embeddings have achieved some success but face limitations in capturing the detailed three-dimensional spatial structure of proteins. To address this issue, researchers have developed representation methods that incorporate not only sequence information but also the geometric features of protein structures.

#### Graph-based Representation Learning

A major advancement in protein representation was the introduction of graph-based learning methods. The three-dimensional structure of proteins is modeled as a graph, where nodes represent atoms or amino acid residues, and edges represent chemical bonds or spatial distances between atoms. Graph Neural Networks (GNNs), such as Graph Convolutional Networks (GCNs) [48] and Graph Attention Networks (GATs) [49], have been employed to learn these representations, enabling the capture of both structural information and interactions between atoms in the protein.

However, standard GNNs lack the ability to maintain equivariance under 3D rotations and translations. This results in learning representations that may be unstable or lack generalization in the context of three-dimensional molecular spaces. To overcome this, SE(3)-equivariant learning methods have been developed to ensure that protein representations remain consistent under geometric transformations.

#### SE(3)-Equivariant Networks

The development of SE(3)-equivariant neural networks marked a significant improvement, ensuring that the learned features of proteins transform correspondingly when subjected to rotations or translations. This is crucial for protein structure-related tasks, as biological interactions occur in three-dimensional space and are independent of specific coordinate systems. Models such as SE(3)-Transformers [50], Tensor Field Networks (TFNs) [51] and Cormorant [36] have been employed to create geometry-aware invariant representations, achieving high performance in tasks like protein-ligand binding prediction.

#### Score-based Generative Models and Diffusion Models

Recently, score-based generative models and diffusion models have proven highly effective in molecular generation. These models, such as Denoising Diffusion Probabilistic Models (DDPMs) [52], employ a stochastic process to gradually add noise to input data, then learn to denoise it in order to recover the original data. This approach is well-suited for molecular structure generation, as molecules and proteins often inhabit complex state spaces with inherent stochasticity in their movements.

### 3.4 Protein ligand generation

Protein-ligand interactions are fundamental to many biological processes, including enzyme catalysis [53], signal transduction [54], and molecular recognition [55]. In the context of drug discovery, these interactions are particularly significant as they form the basis for the development of small-molecule therapeutics. A ligand, typically a small molecule, binds to a target protein, often within a specific region known as the binding pocket, influencing the protein’s function. The goal in drug discovery is to identify ligands that can modulate the activity of disease-related proteins with high specificity and efficacy.

Ligand generation is an essential part of the drug design process, often driven by the need to create novel molecules that exhibit favorable binding properties. Traditional methods rely on screening large compound libraries, which can be time-intensive and computationally expensive. Recent advances in artificial intelligence (AI) and computational chemistry, however, have introduced new paradigms in ligand generation, with AI-driven approaches offering enhanced efficiency and precision.

One of the most promising avenues in ligand generation is the use of generative models. These models, trained on large datasets of known drug-like molecules, can create novel ligands that are optimized for desirable properties such as binding affinity, specificity, and bioavailability. Deep learning methods, particularly variational autoencoders (VAEs) [56], generative adversarial networks (GANs) [57], and diffusion models [52], have been adapted for molecular generation, enabling the design of ligands that fit within the target protein’s binding site or interact favorably with a specific protein conformation.

A key advantage of generative models is their ability to explore chemical space efficiently, proposing novel ligand structures that may not exist in traditional databases. These approaches can generate molecules that meet predefined pharmacophoric requirements, potentially shortening the drug discovery timeline. Moreover, many modern generative approaches, such as structure-based design, incorporate information about the target protein’s 3D structure, leading to ligands that are not only novel but also structurally tailored to interact effectively with the protein of interest.

## 4 Method

The proposed pipeline (as illustrated in Figure 1 and Algorithm 3) is structured into two distinct phases to enhance the efficiency and effectiveness of the drug repurposing process. In the first phase (as illustrated in Algorithm 1), potential binding pockets on the target protein are identified through advanced computational prediction methods. These predicted pockets are then used as specific input conditions for a generative AI model, which is employed to design novel ligand structures optimized for binding to these sites. The generated ligands are treated as initial drug candidates, thereby focusing subsequent screening efforts. In the second phase (as illustrated in Algorithm 2), the pipeline performs similarity searches within existing drug databases, comparing the geometric and chemical properties of the AI-generated ligands to those of approved drugs. This step operates on the foundational assumption that “chemical compounds with similar structures may have similar activities,” a principle that has been a cornerstone of lead identification in drug discovery for many years [10]. Compared to docking simulations, which can be computationally intensive and time-consuming, similarity search-based screening offers a significantly faster alternative, enabling the rapid identification of drug candidates while maintaining high relevance to the target protein.

**Fig. 1:**
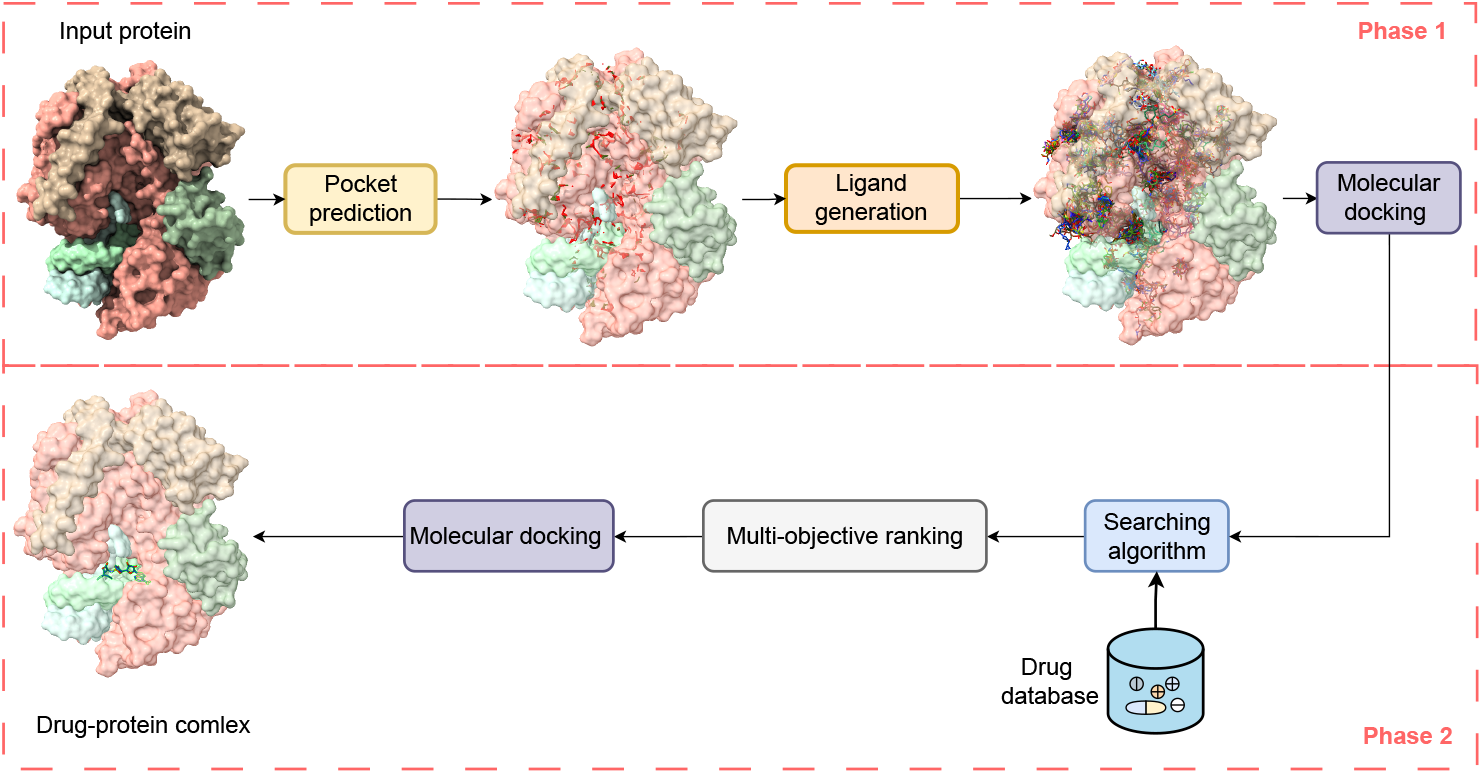
Overview of the Generative AI-assisted Drug Repurposing Pipeline. The pipeline consists of two phases: Phase 1 generates potential ligands using generative AI, and Phase 2 identifies promising drug candidates via similarity-based searches within drug databases.

### 4.1 Phase 1 - Potential ligand generation

#### Pocket prediction

We applied Fpocket[58] as a tool for identifying and characterizing protein pockets, which are potential binding sites for small molecules. It works by detecting cavities on the protein’s surface and then analyzing these cavities to identify important features such as size, volume, and hydrophobicity. These pockets are often crucial for drug design, as they can serve as target sites for inhibitors or other therapeutic compounds.

##### Algorithm 1 Potential ligand generation

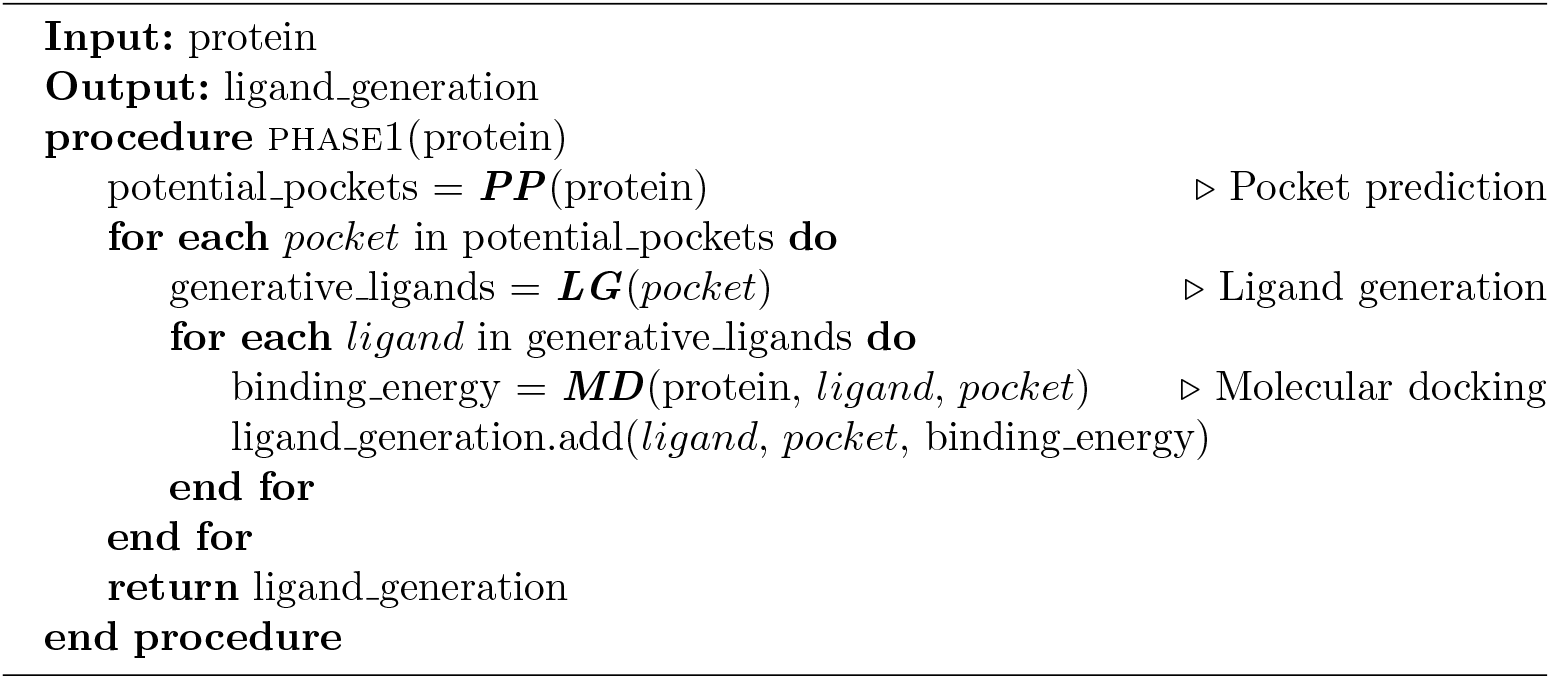

#### Ligand generation

After identifying the potential binding pockets using Fpocket[58], we employed Structure-based Drug Design (SBDD) with Equivariant Diffusion Models [15] to design candidate molecules that can effectively bind to these pockets. SBDD leverages the 3D structure of the protein and its identified pockets to guide the design of ligands that fit optimally into the binding site, improving binding affinity and specificity. The Equivariant Diffusion Models play a critical role by maintaining the geometric and symmetrical properties of molecular structures throughout the generative process, ensuring that the designed molecules conform to the physical constraints of the protein-ligand interactions. These models help generate diverse, high-quality candidate molecules by iteratively refining molecular structures to optimize their fit within the pocket. This approach integrates both the structural information of the protein and the physical properties of potential drug-like molecules, enhancing the efficiency and accuracy of drug discovery efforts.

#### Molecular docking

To calculate the binding energy of generative ligands, we use Vina Docking 3D [59], [60], a molecular docking tool, to simulate and predict how well these ligands interact with the identified binding pockets of the protein. Vina works by exploring possible orientations and conformations of the ligand within the binding site and calculating the binding affinity based on molecular forces such as van der Waals interactions, hydrogen bonding, and electrostatic interactions. By running multiple docking simulations, Vina predicts the most favorable binding pose of the ligand and provides a binding energy score, which reflects the stability and strength of the ligand-protein interaction. This score helps prioritize the best candidates for further optimization or experimental validation, streamlining the process of identifying high-affinity drug candidates in structure-based drug design.

##### Algorithm 2 Potential drug searching

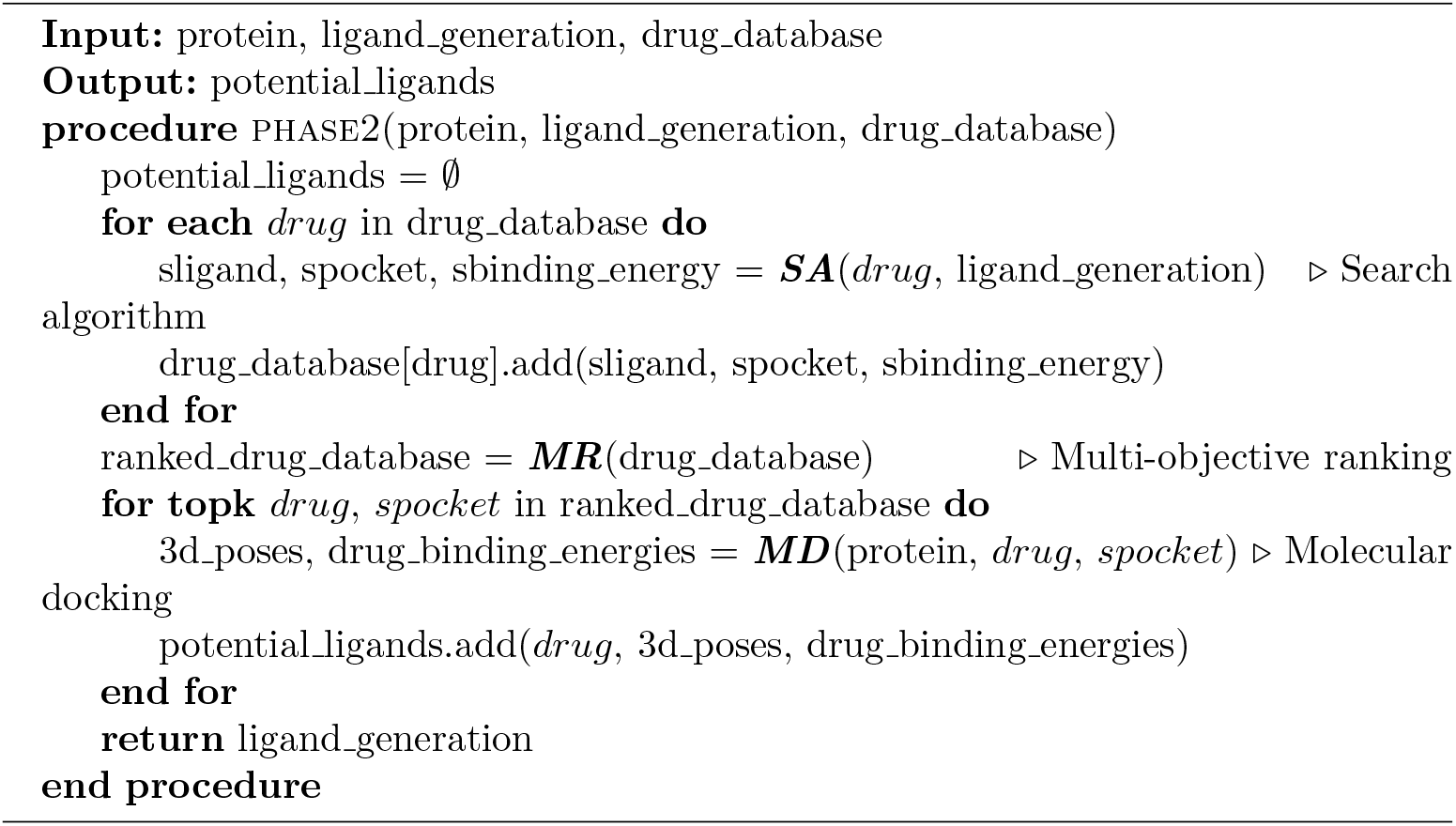

### 4.2 Phase 2 - Potential drug searching

#### Seaching algorithm

We use Deep Graph InfoMax (DGI) [61] global vectors to calculate the similarity between ligands generated from Structure-Based Drug Design (SBDD) and existing compounds in drug databases. DGI is a deep learning technique that extracts meaningful representations from graph-structured data, such as molecular graphs, by maximizing mutual information between global and local graph features. Each molecule is represented as a graph, where nodes correspond to atoms and edges to bonds. By applying DGI, we obtain global vector embeddings for both the generative ligands and the compounds in the drug database. These embeddings capture important structural and chemical features of the molecules. To assess similarity, we compute the cosine similarity between the global vector representations of the generative ligands and the drug database entries. This approach enables us to identify potential analogs or related compounds, facilitating the repurposing of known drugs or suggesting novel molecules with similar pharmacological properties. Additionally, for each drug in the database, we match it with the generative ligand that has the most similar structure to estimate the binding energy and pocket coordinates of the drug.

#### Multi-objective ranking

Rank-Sum Weight Method [62, 63] is used to rank drugs based on both their similarity score and binding energy of generative ligands, ensuring a balanced consideration of multiple criteria. We then sum these ranks to create a final score, optionally assigning different weights to the criteria if one is considered more important. The ligands with the lowest total rank-sum values are prioritized, as they perform well in both binding affinity and structural similarity. This approach enables a robust ranking system that accounts for multiple important factors in drug repurposing pipeline. The fitness function can be formulated as:

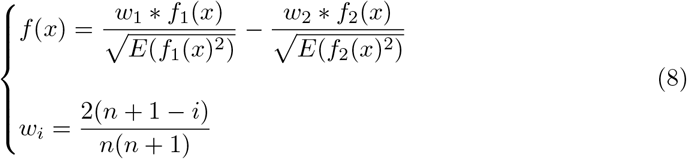

where *x* is a drug in drug database, *f*_1_ is the mapping from a drug in drug database to the similarity score of the most similar ligand in ligand generation database, *f*_2_ is the mapping from a drug database to the binding energy of the most similar ligand in ligand generation database, *E* denotes the expected value (or mathematical expectation) and *n* is number of mappings in the fitness function.

##### Algorithm 3 Generative AI-assisted Drug Repurposing Pipeline

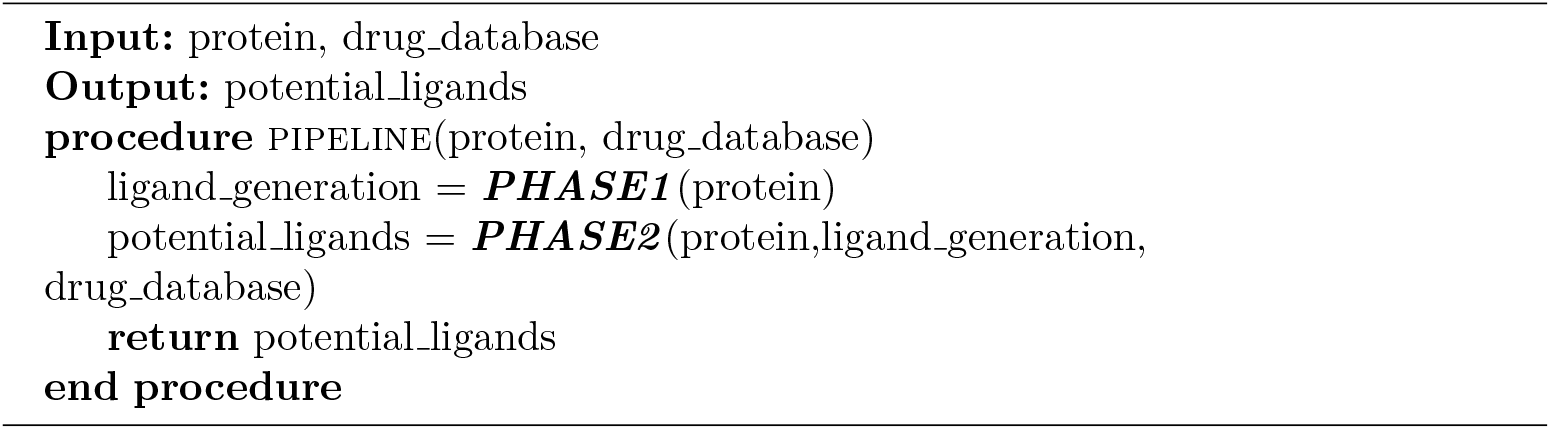

## 5 Experiment

### 5.1 Evaluation on Drug Repurposing Scenarios

To evaluate the generalizability and practical application of our pipeline, we conducted a comprehensive study using the DrugBank drug-target interaction dataset [64], which comprises 846 protein targets and 9716 approved drugs. This evaluation focused on identifying potential drug-target interactions across a broad range of proteins, aiming to verify whether our pipeline could effectively identify real drugs, particularly in drug repurposing scenarios such as COVID-19 and HIV. Given that our screening model relies on binding affinity, we hypothesize that the active compounds identified by the pipeline will have a high probability of being effective drugs in the real world.

However, we acknowledge that not all approved drugs necessarily have the highest binding affinities, as factors beyond affinity, such as pharmacokinetics and safety profiles, also play critical roles in drug approval. To account for this, we decided to report the distribution of the position where the first approved drug for each target is found within our ranked list of predicted compounds.

By analyzing this distribution, we aim to assess how effectively our pipeline can narrow down the search space for wet lab experiments to include the real, approved drugs. Reporting the position of the first approved drug within our ranked list allows us to determine how high these clinically validated drugs appear, even if they do not have the highest predicted binding affinity. Additionally, it is important to consider that the compounds listed above these approved drugs might also represent potential drug candidates that have not yet been tested or validated. This approach not only evaluates the practical utility of the pipeline in reducing the number of compounds that need to be experimentally tested but also highlights the possibility of discovering new, untested compounds that could be viable drug candidates, thereby streamlining the drug discovery process.

### 5.2 Ablation Study

To further assess the performance and efficiency of our Generative AI-assisted Drug Repurposing Pipeline, we conducted an ablation study with two main objectives: (1) to compare the performance of different similarity search methods, and (2) to evaluate the impact of multi-objective ranking on the selection of candidate compounds.

#### Similarity Search Methods

We evaluated the effectiveness of three molecular similarity methods: Tanimoto similarity [65], Morgan fingerprint similarity [66], and Graph Neural Networks (GNNs) [67]. Tanimoto similarity is widely used in cheminformatics to measure the similarity between molecular fingerprints, while Morgan fingerprints encode circular substructures, providing a more detailed representation of molecular architecture. GNNs, as a deep learning-based approach, capture molecular features by modeling molecules as graphs. This comparison aimed to evaluate both the hit rate (the ability to prioritize approved drugs) and computational efficiency (runtime performance) of each method when applied to the DrugBank dataset. Notably, for this ablation study, all similarity search methods were evaluated using the multi-objective ranking approach to ensure a fair comparison across search methods. The goal was to identify which method strikes the best balance between accuracy and computational cost, with hit rate referring to how much the search space is reduced by identifying approved drugs at higher rankings. The hit rate refers to how much the search space is reduced by identifying approved drugs at higher rankings. It is expressed as a percentage reduction in the number of molecules that need to be evaluated, indicating how efficiently each method prioritizes relevant drug candidates. For example, a high hit rate would mean that only a small fraction of the compounds in the database would need further testing, as the method successfully ranks approved drugs near the top of the list. This reduction is crucial, as it significantly decreases the experimental workload, helping to focus resources on fewer compounds that are more likely to be effective. Computational cost, on the other hand, is calculated as the average search time required to search the entire drug database for all target proteins. This metric captures the efficiency of each method in terms of how long it takes to process the data and generate similarity rankings. A method with a lower computational cost would be able to search the database faster, which is particularly important when dealing with large datasets like DrugBank. Balancing a high hit rate with low computational cost is essential, as the most accurate method may not always be the most efficient, and vice versa.

The aim of this comparison was to identify the method that strikes the best balance between accuracy and computational performance.

#### Multi-Objective Ranking

In the second part of the study, we explored the effects of multi-objective ranking on the selection of candidate drugs. We compared single-objective rankings based solely on either similarity scores or binding energy, with a multi-objective approach that combines both criteria. To ensure consistency, we used the GNN-based similarity search method for all ranking strategies, as it demonstrated the best performance in the similarity search ablation study. This analysis focused on assessing the trade-off between hit rate improvement and computational efficiency, specifically docking time, which can be computationally expensive. Docking time, in this context, refers to the average time required to computationally dock a drug candidate to its target protein, starting from the highest-ranked compound down to the first approved drug in the database. Docking is a critical step in drug discovery, as it simulates the binding of small molecules (drug candidates) to target proteins to evaluate their potential efficacy. The docking process, however, can be highly computationally intensive, especially when applied to large datasets or when high accuracy is required in the simulations. In our study, docking time was used as a measure of computational cost, representing how long it takes to perform detailed binding simulations for candidate drugs. A key focus was to determine whether the multi-objective ranking approach—considering both similarity scores and binding energy—could reduce the number of docking simulations required to identify approved drugs. By narrowing down the search space through similarity-based rankings, the goal was to prioritize higher-quality candidates and minimize docking time by focusing on fewer, more promising molecules.This part of the study was conducted on a smaller subset of the DrugBank dataset, carefully sampled to retain the overall distribution of the original dataset, in order to mitigate the computational cost. Ultimately, the aim was to evaluate whether combining multiple objectives in the ranking process could improve both the efficiency of docking and the accuracy of selecting approved drugs, providing a more balanced approach without a significant increase in computational burden.

### 5.3 Results and Discussion

This section evaluates the performance of our Generative AI-assisted Drug Repurposing Pipeline, with a focus on its ablation study and drug repurposing results. For further context, detailed descriptions of the metrics used are provided in Appendix A, the computational resources utilized are outlined in Appendix B, and additional results supporting our findings are presented in Appendix C.

#### 5.3.1 Ablation Study Results

In the similarity search evaluation on the DrugBank DTI dataset, we assessed the efficacy of different methods in identifying approved drugs, as summarized in Table 2. The results indicate that GNN-based approaches outperformed traditional similarity measures like Tanimoto and Morgan fingerprints, as well as advanced graph-based techniques such as GATs, EGNNs, and Equiformer. Specifically, EGNNs achieved the highest hit rate reduction of 60.67%, demonstrating a notable improvement in prioritizing relevant drug-target interactions. However, GNNs distinguished themselves by balancing high hit rate reduction (55.76%) with the shortest average runtime of only 12.95 seconds, significantly outperforming traditional methods, which exhibited much longer runtimes (714.40 seconds for Tanimoto and 786.98 seconds for Morgan fingerprints). Other graph-based models like GATs and Equiformer delivered competitive hit rate reductions of 58.36% and 57.31%, respectively, but with moderately higher runtimes. These findings emphasize the computational efficiency of GNNs, which deliver robust performance gains in hit rate reduction and runtime, positioning them as a highly effective solution for large-scale drug similarity searches. In the multi-objective ranking study, we assessed the impact of combining similarity scores and binding energy (SS + BE) as compared to using either criterion alone. As detailed in Table 3, the multi-objective approach yielded the best performance, with a hit rate reduction of 61.24%, outperforming both single-objective methods. While the SS + BE approach increased the docking time to 49.97 seconds, this is a reasonable trade-off given the significant improvement in ranking accuracy. The single-objective ranking based on binding energy (BE) alone was the fastest method, with an average docking time of 48.65 seconds, but it came at the cost of reduced hit rate performance (55.74%). This demonstrates that the multi-objective ranking strategy provides a balanced approach, optimizing both hit rate and computational efficiency in the drug repurposing pipeline.

**Table 2:** Comparison of Similarity Search Methods on the DrugBank DTI Dataset.

| Search Method | Hit Rate Reduction (%) | Runtime (s) |
| --- | --- | --- |
| Tanimoto | 55.28 | 714.40 |
| Morgan Fingerprint | 58.12 | 786.98 |
| GNNs | 55.76 | **12.95** |
| GATs | 58.36 | 13.86 |
| EGNNs | **60.67** | 50.77 |
| Equiformer | 57.31 | 83.24 |

**Table 3:** Comparison of Single-Objective and Multi-Objective Ranking Methods on the DrugBank DTI Dataset.

| Ranking Method | Hit Rate Reduction (%) | Docking Time (s) |
| --- | --- | --- |
| Similarity Score (SS) | 59.33 | 52.61 |
| Binding Energy (BE) | 55.74 | **48.65** |
| SS + BE | **61.24** | 49.97 |

#### 5.3.2 Drug Repurposing Evaluation Results

In this section, we closely examine the results of our Generative AI-assisted Drug Repurposing Pipeline, configured with the optimal combination of GNN-based similarity search and multi-objective ranking, as established in our ablation study. This analysis specifically highlights the effectiveness of this setup on the DrugBank drug-target interaction dataset. As depicted in Figure 2, our approach yields a strong concentration of approved drugs in the higher-ranked positions. Notably, the median position of the first approved drug per target protein is approximately 2000, meaning that half of these drugs appear within the top 2000 ranks. Furthermore, the lower quartile (Q1) is near 1000, and the upper quartile (Q3) around 4000, with only a few outliers reaching the maximum rank of 9716. This distribution illustrates that the combination of GNN-based search and multi-objective ranking significantly enhances the prioritization of therapeutically relevant compounds, effectively narrowing down the search space and making the discovery of potential drug candidates more efficient. To illustrate, we provide some examples from the DrugBank dataset. The pipeline ranked Acarbose, an inhibitor of human pancreatic alpha-amylase (HPA) [68], at position 253 out of 9,716, placing it within the top 3%, and Cefoperazone, a beta-lactam antibiotic targeting penicillin-binding protein 1a (PBP1a) [69], at position 52 out of

**Fig. 2:**
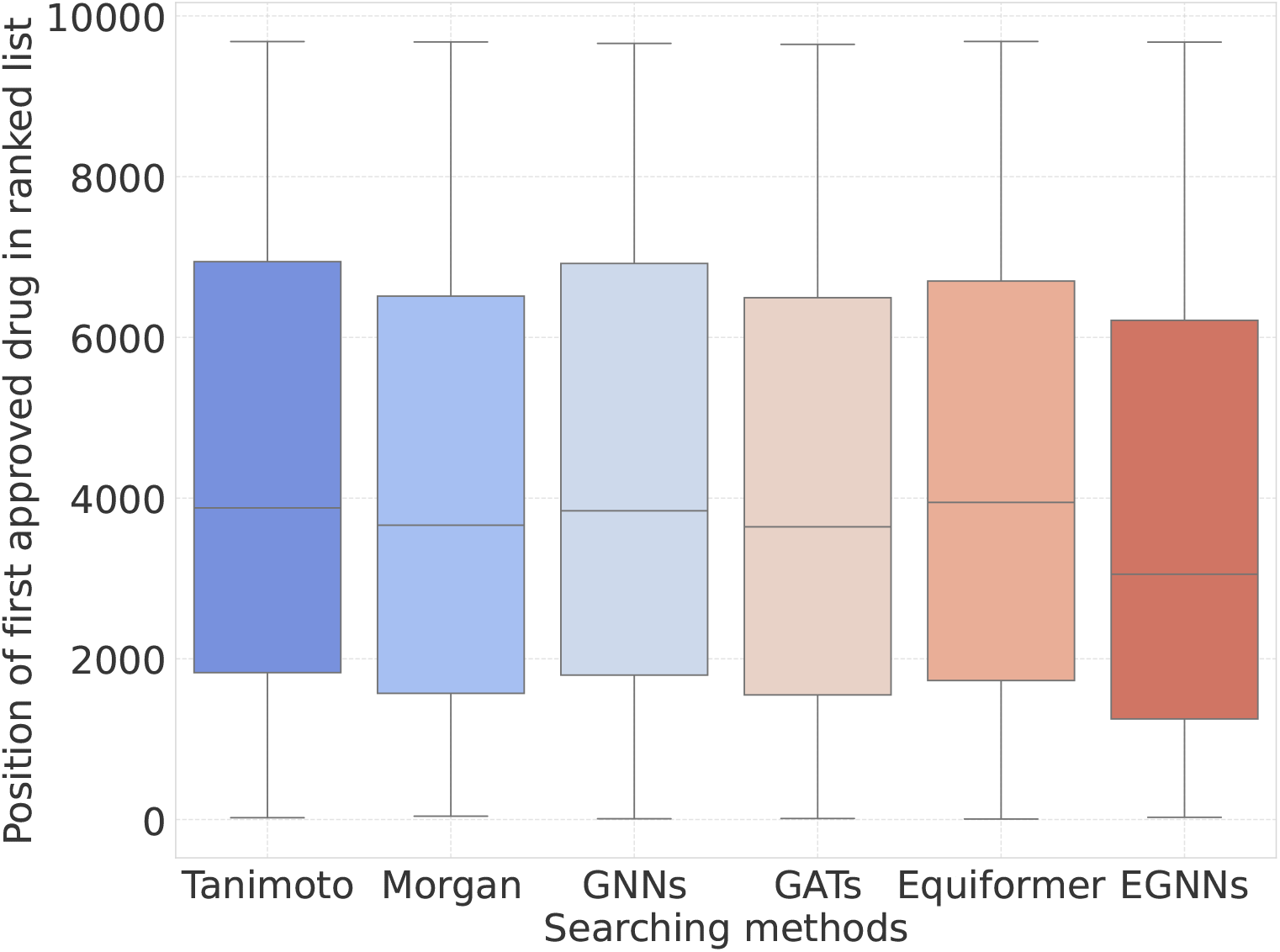
Distribution of the rank positions of the first approved drug for each protein target in the DrugBank DTI dataset across different searching methods.

9,716, within the top 1%. Additionally, in our case studies, the pipeline performed exceptionally well for specific targets such as HIV and COVID-19. For HIV, the pipeline ranked Abacavir, a well-established antiretroviral drug used in combination therapy to inhibit reverse transcriptase [70], at position 1,055 out of 9,716, placing it within the top 10% of candidates. In the COVID-19 case study, the pipeline identified Remdesivir, an antiviral targeting the RNA-dependent RNA polymerase (RdRp) of SARS-CoV-2 to inhibit viral replication [71], at position 637 out of 9,716, within the top 7%.

Beyond generating a ranked list of potential drugs for each target, our pipeline also provides detailed drug-target complex conformations, offering valuable insights into the binding interactions at the molecular level. As illustrated in Figure 3, the pipeline effectively predicts drug poses that closely align with experimentally determined structures, even without prior information about specific binding pockets. In this figure, the predicted drug conformations are overlaid with the experimentally validated poses, showing a strong correspondence between the predicted and actual binding modes. This high degree of alignment demonstrates the pipeline’s ability to accurately model drug-target interactions solely from 3D structural data, further highlighting its robustness and effectiveness.

**Fig. 3:**
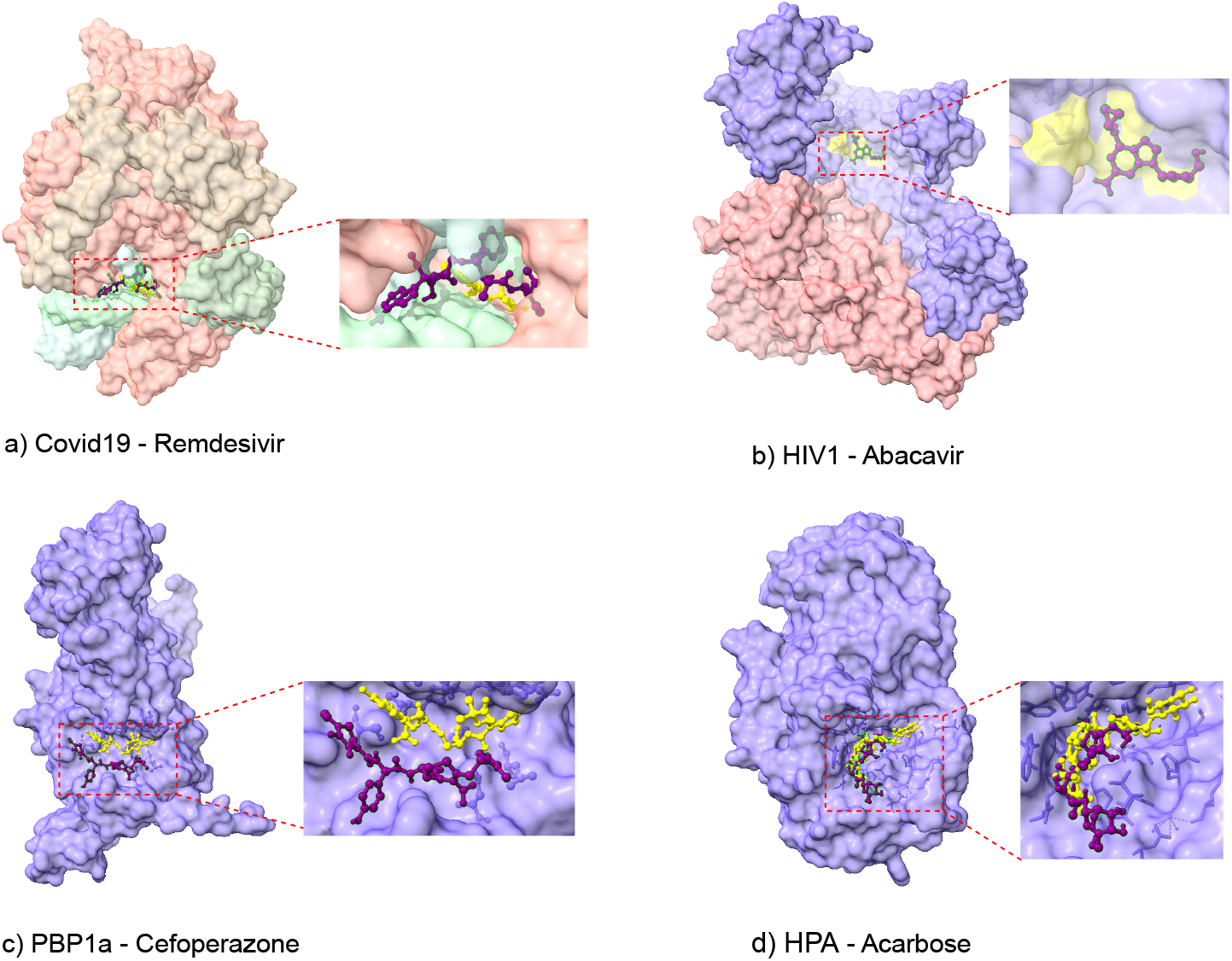
Comparison of predicted drug-target complexes by our pipeline with experimentally determined structures: (a) COVID-19 - Remdesivir, (b) HIV-1 - Abacavir, (c) PBP1a - Cefoperazone, (d) HPA - Acarbose. The predicted drug poses are shown in purple, while the experimentally determined drug poses are shown in yellow, demonstrating the alignment between the predicted and real drug conformations. For the case of HIV-1 and Abacavir (b), where the 3D complex structure is unavailable, the true binding site is highlighted in yellow to indicate the experimentally validated pocket region.

#### 5.3.3 Discussion

The conceptual framework of the pipeline is intentionally expansive, structured to incorporate a diverse array of modular components, each of which plays a critical role in rigorously validating distinct facets of the overarching concept. The selection of these modules is neither arbitrary nor incidental; rather, it is informed by a deliberate strategy aimed at ensuring the modules’ capacity to verify discrete stages within the pipeline, thereby safeguarding both flexibility and structural integrity. This methodical approach facilitates a comprehensive and incremental validation process, as evidenced by the detailed findings presented in Section 5.3.1 and Section 5.3.2. Furthermore, the model is intrinsically adaptable, designed to effortlessly assimilate state-of-the-art (SOTA) modules. By capitalizing on advancements within the specific scientific domains from which these modules are derived, the model is positioned to continuously evolve and enhance its performance, remaining future-ready to integrate cutting-edge innovations as research in these fields progresses.

## 6 Conclusion

This paper introduces a Generative AI-assisted Virtual Screening Pipeline designed to enhance the efficiency and generalizability of drug repurposing. By leveraging generative AI models for ligand generation, binding pocket prediction, and similarity-based searches within approved drug databases, our pipeline significantly accelerates the identification of new therapeutic uses for existing drugs. The results from our experiments on the DrugBank dataset, particularly in the case studies of HIV and COVID-19, highlight the pipeline’s potential to efficiently prioritize promising drug candidates. Notably, these achievements were realized without requiring any prior knowledge of the target proteins or drugs beyond their 3D structures and without relying on training with drug-target interaction datasets. This capability underscores the pipeline’s robustness and adaptability, making it particularly valuable for addressing emerging health threats and novel targets where data is scarce or unavailable. Future work will focus on developing additional specialized pipelines to further enhance the generalizability and effectiveness of our drug repurposing strategies, ensuring broader applicability across diverse therapeutic contexts.

## A Metrics Description

This section provides a detailed explanation of the metrics used to evaluate the performance of similarity search methods and ranking strategies in our study.

### Hit Rate Reduction (%)

Hit rate reduction measures the proportion of the search space eliminated by prioritizing approved drugs within higher ranks in the list of compounds. It is calculated as:

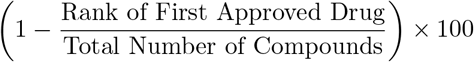

The reported value represents the average hit rate reduction across all drug-target pairs in the dataset. Higher values indicate better prioritization of approved drugs, effectively narrowing down the search space and improving computational efficiency.

### Runtime (s)

This metric represents the average time required for a similarity search method to compute all similarity scores for a single protein target across the entire dataset of molecules. Specifically, it measures the total time taken for embedding generation and pairwise similarity computations for one protein target. The reported value is the mean runtime across all protein targets in the dataset.

### Docking Time (s)

Docking time measures the average time needed to perform docking simulations to identify potential drugs. For each protein target, this metric calculates the average docking time required to process molecules from the top of the ranked list down to the first approved drug (first hit). This value is then averaged across all protein targets in the dataset. The purpose of this metric is to evaluate how effectively a similarity search method prioritizes suitable molecules for docking. Since docking simulations for unsuitable molecules often take significantly longer, this metric highlights the ability of our method to rank potential drugs higher, thereby reducing computational costs and accelerating the discovery process.

These metrics collectively provide a comprehensive evaluation framework, balancing accuracy (hit rate reduction) and computational efficiency (runtime and docking time) to assess the suitability of various methods in the drug repurposing pipeline.

## B Computational Resources

All experiments in this study were conducted using NVIDIA A100 GPUs, which provided the computational power necessary for handling large-scale datasets and executing complex similarity search tasks efficiently. Each A100 GPU is equipped with 40 GB of memory, supporting the seamless processing of deep learning models and rapid data throughput. This high-performance setup ensured consistent and optimal conditions for comparing computational efficiency across different approaches. Additionally, all time measurements, including docking and similarity search runtimes, were obtained under this configuration to ensure standardized performance assessments.

## C Additional Results

This section aims to provide further insights into the performance of various similarity search methods tested on the DrugBank DTI dataset. As illustrated in Figure 4, the distribution of protein targets based on the ranking position of their first approved drug across different methods reveals distinct patterns of effectiveness. Morgan Fingerprints demonstrate a slightly better focus on higher-ranked positions compared to GNNs and Equiformer, aligning with their strong performance in prioritizing approved drugs. EGNNs display the sharpest concentration of approved drugs in top-ranked positions, confirming their superior ability to identify relevant compounds. Meanwhile, GATs exhibit a balanced distribution, further highlighting their consistency across ranking tasks.

**Fig. 4:**
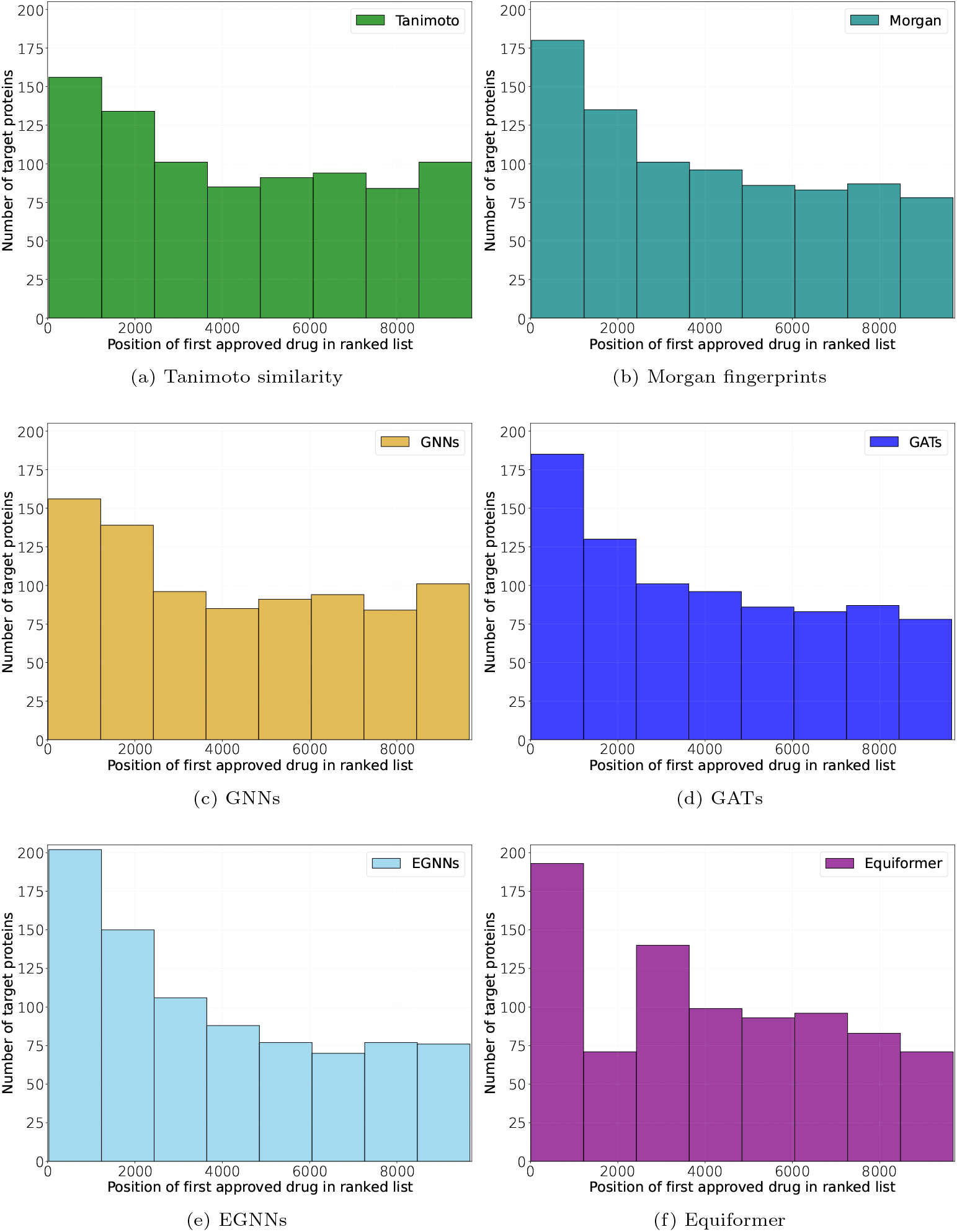
Distribution of protein targets based on the ranking position of their first approved drug in the DrugBank DTI dataset. The x-axis represents the rank position of the first approved drug (ranging from 0 to 9717), while the y-axis shows the number of protein targets associated with each rank. This visualization highlights the effectiveness of various searching methods in prioritizing approved drugs across the dataset.

In addition to ranking positions, we evaluated the ADMET properties [72] (Absorption, Distribution, Metabolism, Excretion, and Toxicity) of drugs for the ranked lists generated in two case studies: HIV and COVID-19. ADMET properties are critical for assessing drug potential, as they provide insights into a compound’s pharmacokinetic and safety profile, which are essential for evaluating its suitability as a therapeutic candidate. Due to the size of the resulting tables, detailed results are available in our public repository where readers can explore the complete data: https://github.com/HySonLab/DrugPipe.

